# Repurposing CD19-directed immunotherapies for pediatric t(8;21) acute myeloid leukemia

**DOI:** 10.1101/2024.04.19.590200

**Authors:** Farnaz Barneh, Joost B. Koedijk, Noa E. Wijnen, Tom Meulendijks, Minoo Ashtiani, Ester Dunnebach, Noël Dautzenberg, Annelisa M. Cornel, Anja Krippner-Heidenreich, Kim Klein, C. Michel Zwaan, Jürgen Kuball, Stefan Nierkens, Jacqueline Cloos, Gertjan J.L. Kaspers, Olaf Heidenreich

**Affiliations:** Princess Máxima Center for Pediatric Oncology, Utrecht, The Netherlands; Erasmus MC-Sophia Children’s Hospital, Department of Pediatric Oncology, Rotterdam, The Netherlands; Emma Children’s Hospital, Amsterdam UMC, Vrije Universiteit, Department of Pediatric Oncology, Amsterdam, The Netherlands; Center for Translational Immunology, University Medical Center Utrecht, Utrecht University, Utrecht, The Netherlands; Wilhelmina Children’s Hospital/University Medical Center, Utrecht, The Netherlands; Department of Hematology, Amsterdam UMC, Vrije Universiteit Amsterdam, Cancer Center Amsterdam, The Netherlands; Wolfson Childhood Cancer Research Centre, Translational and Clinical Research Institute, Newcastle University, Newcastle upon Tyne, United Kingdom; University Medical Center Utrecht, Utrecht University, Utrecht, The Netherlands

**Keywords:** children, tumor immune microenvironment, blinatumomab, CAR T cells, tumor antigens

## Abstract

In contrast to patients with B cell precursor acute lymphoblastic leukemia (BCP-ALL), patients with acute myeloid leukemia (AML) have not yet benefited from recent advances in targeted immunotherapy. Repurposing immunotherapies that have been successfully used to target other hematological malignancies could, in case of a shared target antigen, represent a promising opportunity to expand the immunotherapeutic options for AML. Here, we evaluated the expression of CD19 in a large pediatric AML cohort, assessed the *ex vivo* AML killing efficacy of CD19-directed immunotherapies, and characterized the bone marrow immune microenvironment in pediatric AML, BCP-ALL, and non-leukemic controls. Out of 167 newly diagnosed *de novo* pediatric AML patients, 18 patients (11%) had CD19^+^ AML, with 61% carrying the translocation t(8;21)(q22;q22). Among CD19^+^ samples, we observed a continuum of CD19 expression levels on AML cells. In individuals exhibiting unimodal and high CD19 expression, the antigen was consistently present on nearly all CD34^+^CD38^-^ and CD34^+^CD38^+^ subpopulations. In *ex vivo* AML-T cell co-cultures, blinatumomab demonstrated substantial AML killing, with an efficacy similar to BCP-ALL. In addition, CAR T cells could effectively eliminate CD19^+^ AML cells *ex vivo*. Furthermore, our immunogenomic assessment of the bone marrow immune microenvironment of newly diagnosed pediatric t(8;21) AML revealed that T- and NK cells had a less exhausted and senescent phenotype in comparison to pediatric BCP-ALL. Altogether, our study underscores the promise of CD19-directed immunotherapies for the treatment of pediatric CD19^+^ AML.

## Introduction

Although the survival of children with acute myeloid leukemia (AML) has improved considerably over the past decades, 25-35% of patients face relapse, which still has an unfavorable prognosis.^1–3^ In addition, current high-dose chemotherapy and allogeneic stem cell transplantation (allo-SCT) regimens lead to significant side and late effects, together illustrating the need for more effective and less toxic therapeutic options.^3^ Nontransplant, targeted immunotherapies such as bispecific antibodies and CAR T cells are of interest given their successes in both solid and hematological malignancies.^4–6^ However, the development of targeted immunotherapy for AML has been challenging, mainly due to the paucity of tumor-specific antigens, on-target off-leukemia hematotoxicity when targeting myeloid-lineage antigens, and the immunosuppressive tumor microenvironment.^7, 8^ Accordingly, with the exception of the CD33-directed antibody-drug-conjugate gemtuzumab ozogamicin, no targeted immunotherapeutic agents have been approved for adults or children with AML.^9, 10^ Hence, repurposing immunotherapies that have been successfully used to target other hematological malignancies could, in case of a shared target antigen, represent a promising opportunity to expand the immunotherapeutic options for AML.

CD19 is a B cell marker which is highly expressed on B cell precursor acute lymphoblastic leukemia (BCP-ALL) cells. For children and adults with BCP-ALL, the CD19-directed bispecific T cell-engager blinatumomab (CD3 x CD19) and CD19-directed chimeric antigen receptor (CAR) T cells (tisagenlecleucel) demonstrated promising results in both pediatric and adult BCP-ALL, which ultimately led to their clinical approval.^6, 11–15^ In AML, expression of CD19 is characteristic for t(8;21)(q22;q22), the most common translocation in children with this disease.^16^ Interestingly, two case studies have reported complete molecular responses in two adults with relapsed CD19^+^ t(8;21) AML following treatment with either blinatumomab or CD19-directed CAR T cells.^17, 18^ Furthermore, two clinical trials are currently investigating the efficacy of CD19-directed CAR T cells in relapsed and refractory (R/R) adult AML (NCT04257175 and NCT03896854). In several pediatric hematological malignancies including AML, another clinical trial is testing a combination of T cell-directed immunotherapies including blinatumomab in R/R CD19^+^ patients after allo-SCT (NCT02790515). Apart from these case studies and ongoing trials, CD19-directed immunotherapies have not yet been studied in pediatric or adult AML and therefore, its efficacy in AML remains unknown.

Here, we examined the expression of CD19 on AML cells in a large cohort of children with newly diagnosed *de novo* AML, evaluated the *ex vivo* efficacy of CD19-directed immunotherapies, and characterized the bone marrow (BM) immune microenvironment in pediatric AML, BCP-ALL, and non-leukemic controls. Our work reveals pediatric t(8;21) AML as a subgroup with a high percentage of CD19^+^ patients, and CD19^+^ t(8;21) AML to be sensitive to blinatumomab- and CAR T cell-mediated cytotoxicity. Furthermore, our immunogenomic analyses of the BM immune microenvironment show that T- and NK cells in pediatric t(8;21) AML have a less exhausted and senescent phenotype in comparison to pediatric BCP-ALL. Altogether, our study demonstrates the potential of CD19-directed immunotherapies for the treatment of pediatric CD19^+^ t(8;21) AML.

## Materials and Methods

### Ethical regulations

This study complied with all relevant ethical regulations and was approved by the Institutional Review Board of the Princess Máxima Center (PMCLAB2021.207, PMCLAB2021.258, and PMCLAB2022.334).

### Clinical and flow cytometry data

A retrospective medical records analysis identified pediatric patients with newly diagnosed *de novo* AML treated in Dutch hospitals between January 2012 and October 2022 (details on treatment regimen and response definitions are provided in Supplementary Methods). Reports of flow cytometry data collected in the diagnostic and, if applicable, relapse setting were retrieved to screen patients for CD19 positivity according to the guideline for assessment of marker positivity by the Dutch Foundation for Quality Assessment in Medical Laboratories (SKML; Supplementary Methods).^19^

### CD19 expression analysis

The flow cytometry standard files, which were accessible for seven t(8;21) AML patients in the total study cohort, were utilized to examine the CD19 expression on myeloid blasts. Flow cytometry results were analyzed using FlowJo™ (v10.10 Software; BD Life Sciences) or Kaluza Analysis software (v2.2.1.20183; Beckman Coulter). The gating strategy for defining AML blasts is shown in **Figure 3A**. The coverage of CD19 expression among AML subpopulations was further determined by analyzing BM mononuclear cells (MCs) from two t(8;21) AML patients and one t(8;21) AML patient-derived xenograft (PDX) sample (RL048)^20^ by flow cytometry (Cytoflex LX, Beckman Coulter; antibodies provided in Supplementary Methods).

### *Ex vivo* T cell killing assays

For allogeneic killing assays, healthy donor CD3^+^ T cells isolated from healthy donors were first expanded using a previously published rapid expansion protocol.^21^ Subsequently, T cells were co-cultured with CD19^+^ primary t(8;21) AML BMMCs, RL048 PDX cells, or primary BCP-ALL BMMCs, at various effector-to-target (E:T) ratios (Supplementary Methods). Next, blinatumomab (1 nM; Blincyto^®^, Amgen) was added to the co-cultures for 48 hours, and co-cultures in the absence of blinatumomab were used to determine the extent of background killing. In autologous killing assays with primary BMMCs, blinatumomab (1 nM) was added directly to unsorted samples (1 x 10^5^ BMMCs per well on a 96-well plate) to activate T cells present in the sample. The viability of blinatumomab-treated samples was normalized to conditions without blinatumomab. Details on the T cell activation and Interferon-γ secretion procedures are provided in Supplementary Methods.

### Bulk RNA-sequencing

Bulk RNA-sequencing (RNA-seq) and data analysis was performed according to the institute’s standard pipelines.^22, 23^ Details on the comparison of gene expression profiles between CD19^+^- and CD19^-^ t(8;21) AML are provided in Supplementary Methods. To characterize the BM immune microenvironment of patients with CD19^+^- and CD19^-^ t(8;21) AML (n=10), AML with other genotypes (n=30; cytogenetic data and other clinical parameters are shown in **Supplementary Table 1**), BCP-ALL (n=209; cytogenetic data and other clinical parameters are depicted in **Supplementary Table 2**), and non-leukemic controls (n=4)^26^, we applied the immune deconvolution platform CIBERSORTx (cibersortx.stanford.edu; LM22 reference signature) to estimate the abundance of lymphoid populations and the TIDE algorithm to infer the abundance of several myeloid and stromal cell types (tide.dfci.harvard.edu; rationale for selected cell populations and details of other immune-based scores are provided in Supplementary Methods).^27–30^

### Statistical analyses

All data were analyzed using the SPSS software (v26.0.0.1; IBM, USA) and GraphPad Prism (v8.0.2; GraphPad Software, USA). Two-sided *P* values of < 0.05 were considered statistically significant. Details of statistical methods and tests are provided in Supplementary Methods.

## Results

### CD19 expression is enriched in pediatric t(8;21) AML

To investigate CD19 as a potential target antigen in pediatric AML, we examined 167 *de novo* pediatric AML patients for CD19 positivity at diagnosis and, when applicable, at relapse. Using records of diagnostic flow cytometry data, we identified 18 newly diagnosed patients (11%) with CD19^+^ AML cells (n=8 with CD19 median fluorescence intensity (MFI) difference (Δ) between leukemic blasts and healthy population of >10-fold, n=10 with ΔMFI of 3 to 10-fold; **Figure 1A)**. We next explored whether CD19 expression was associated with specific cytogenetic alterations. In line with data in adult AML, we found that 61% (n=11) of CD19^+^ patients carried the translocation t(8;21)(q22;q22) (n=3: CD19 ΔMFI >10-fold, n=8: 3 to 10-fold; **Figure 1B**).^33, 34^ Other cytogenetic alterations of CD19^+^ pediatric AML patients included t(9;11)(p22;q23) (n=2), t(16;21)(q24;q22) (n=2), t(1;11)(q21;q23) (n=1), inv(16)(p13;q22) (n=1), and in one case cytogenetic information was not available (**Figure 1B**; additional clinical characteristics are listed in **Table 1**). In the entire cohort, 21 out of 167 patients had the (8;21) translocation, indicating that 52% of patients with this translocation were CD19^+^ (**Figure 1C)**.

**Figure 1.**
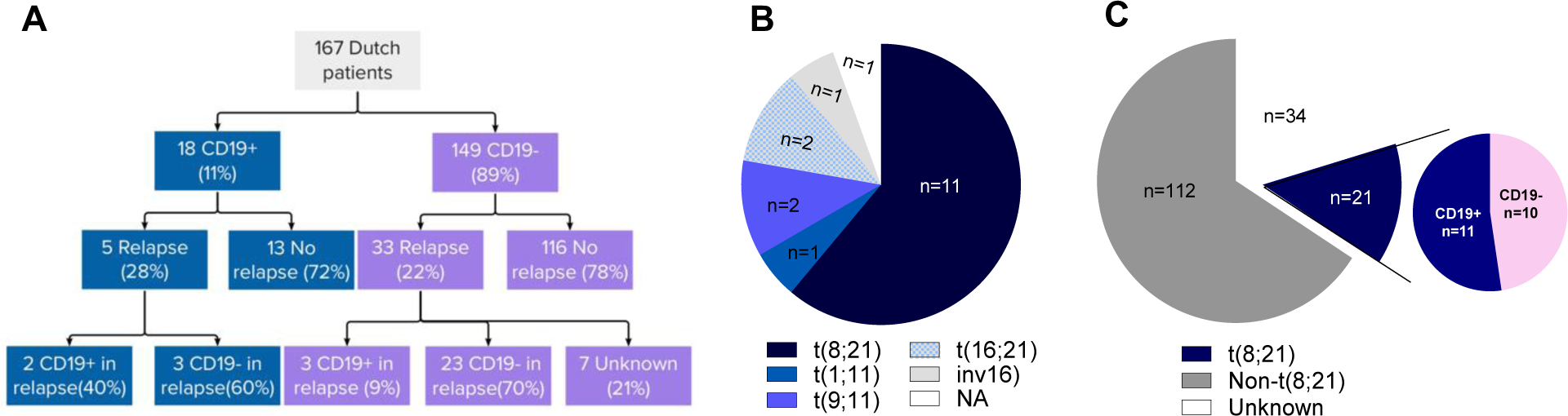
CD19 expression among pediatric AML patients. (A) Incidence of CD19 positivity among newly diagnosed and relapsed pediatric AML patients. (B) Cytogenetic alterations observed in CD19^+^ pediatric AML patients. NA=not available. (C) Incidence of the t(8;21) subtype across the total cohort, and the incidence of CD19 positivity among all t(8;21) patients.

**Table 1.**
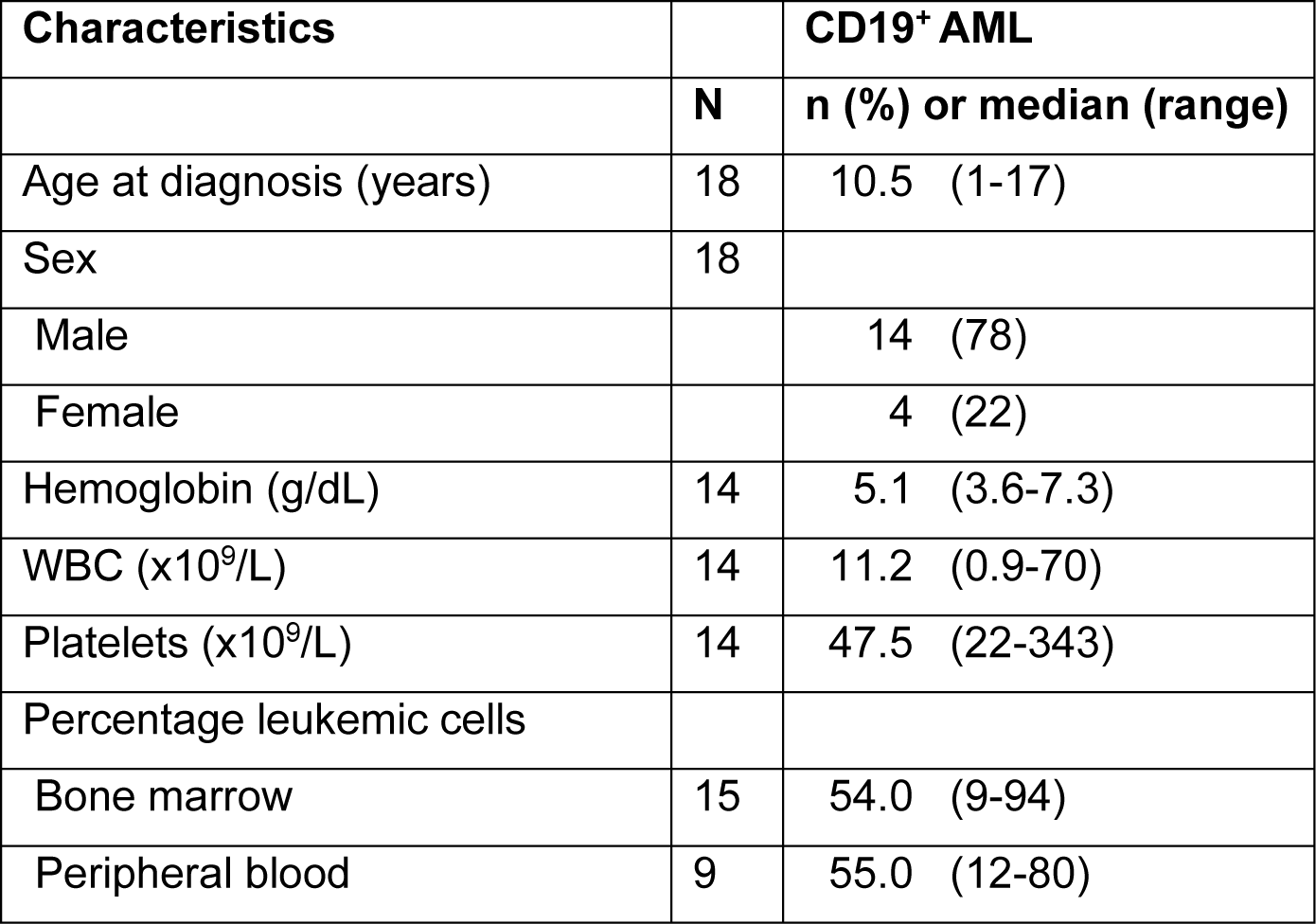
Baseline characteristics of CD19^+^ pediatric AML patients at diagnosis.

| Characteristics |  | CD19<sup>+</sup> AML |
| --- | --- | --- |
|  | N | n (%) or median (range) |
| Age at diagnosis (years) | 18 | 10.5 (1-17) |
| Sex | 18 |  |
| Male |  | 14 (78) |
| Female |  | 4 (22) |
| Hemoglobin (g/dL) | 14 | 5.1 (3.6-7.3) |
| WBC (x10<sup>9</sup>/L) | 14 | 11.2 (0.9-70) |
| Platelets (x10<sup>9</sup>/L) | 14 | 47.5 (22-343) |
| Percentage leukemic cells |  |  |
| Bone marrow | 15 | 54.0 (9-94) |
| Peripheral blood | 9 | 55.0 (12-80) |

Regarding patient outcomes of CD19^+^ AML patients, all (n*=*18) achieved complete remission (CR) by the end of the second induction course (100%), and there were no early deaths. Among the patients with CD19^+^ AML at diagnosis, five experienced disease relapse (**Figure 1A**). In two of these cases, AML cells retained CD19 positivity, and both patients are currently alive. Conversely, in three cases, AML cells lost their CD19 expression at relapse. One of these patients deceased, which was the only death among patients with CD19^+^ AML at diagnosis. Intriguingly, three patients with CD19^-^ AML at diagnosis gained CD19 positivity at relapse (out of 33 relapses in the CD19^-^ AML group), with a fatal outcome in one of these patients.

Next, we investigated whether CD19 expression on AML cells was associated with event-free survival (EFS) and overall survival (OS). To account for the confounding effect of cytogenetic alterations, we compared EFS and OS among all t(8;21) patients (n=21). In this exploratory analysis, the 5-year EFS and 5-year OS for the CD19^+^ (n=11) and CD19^-^ (n=10) groups were 65% (SE 17) and 74% (SE 16), and 100% and 89% (SE 11), respectively, showing no substantial difference, although the cohort size was limited (**Figure S1A-B**). Taken together, our data reveal an enrichment of CD19 positivity in pediatric t(8;21) AML at diagnosis, and gain of CD19 expression in relapsed cases with CD19^-^ disease at initial diagnosis.

### CD19^+^ t(8;21) AML exhibits reduced metabolic activity and cell division

We next sought to further characterize differences between CD19^+^ and CD19^-^ t(8;21) AML. Specifically, given the typical expression of B cell-related genes such as *CD19* and *PAX5*, a B lymphoid transcription factor responsible for *CD19* upregulation, in t(8;21) AML, we aimed to investigate whether gene expression programs seen in (pre-)B cells were present in CD19^+^ t(8;21) AML. To investigate this, we retrieved BM bulk RNA-sequencing data of patients with CD19^+^- and CD19^-^ t(8;21) AML and a blast percentage of at least 75% (n=6 vs. n=3, respectively). As anticipated, differential gene expression analysis revealed significant upregulation of the B cell-related genes *CD19* and *POU2AF1* in CD19^+^ t(8;21) AML, as well as a trend towards higher expression of *PAX5 (***Figure 2A-B** and **Supplementary Table 3**).^35^ Furthermore, GSEA showed that CD19^+^ t(8;21) AML demonstrated a decrease in various metabolic processes including oxidative phosphorylation and fatty acid metabolism in comparison to CD19^-^ t(8;21) AML, suggestive of lower metabolic activity in CD19^+^ t(8;21) AML (**Figure 2C** and **Figure S2**). These data are consistent with previous work showing that *PAX5* enforces a state of chronic energy deprivation in pre-B cells.^36^ In addition, cell cycle-related pathways were depleted in CD19^+^-compared to CD19^-^ t(8;21) AML, together suggesting a less proliferative state in CD19^+^ t(8;21) AML (**Figure 2C** and **Figure S2**). Given that such cells show in general less susceptibility to conventional chemotherapy, these data suggest that alternative therapies such as immunotherapies could be a suitable treatment option for CD19^+^ AML.^37^

**Figure 2.**
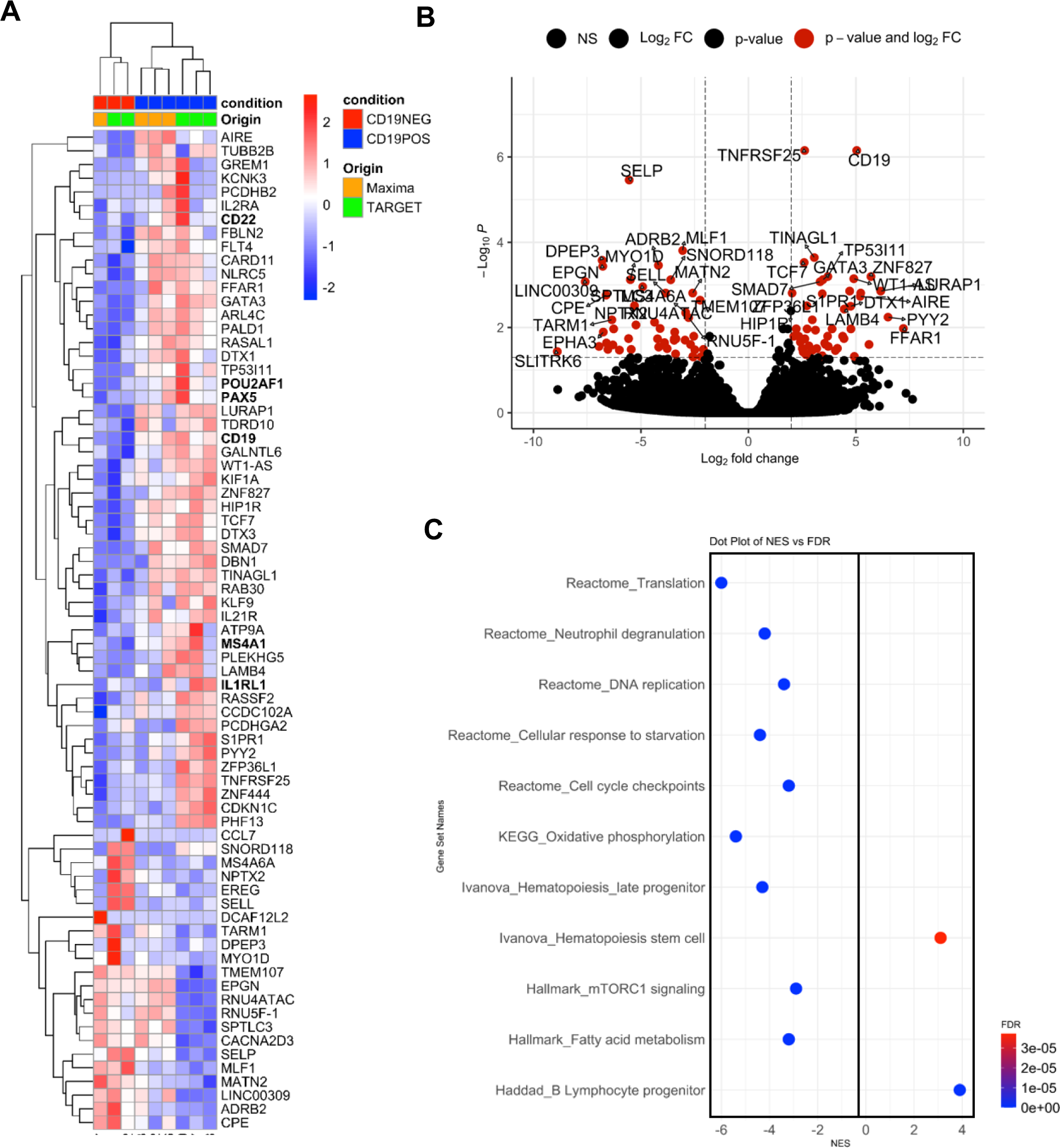
Transcriptomic differences between CD19^+^ and CD19^-^ t(8;21) AML. (A) Heatmap showing the expression of the top up- and downregulated genes between CD19^+^-(n=6) and CD19^-^ (n=3) t(8;21) AML for each patient. The color bar indicates the logarithmically scaled and normalized gene expression values. (B) Volcano plot showing the differentially expressed genes between CD19^+^-(n=6) and CD19^-^ t(8;21) AML. (C) Gene set enrichment analysis plot showing enriched and depleted phenotypes and pathways in CD19^+^-compared to CD19^-^ t(8;21) AML. FDR <0.05 was considered significant. NES: normalized enrichment score. FDR: false discovery rate.

### CD19 is expressed among different t(8;21) AML subpopulations

To evaluate the suitability of CD19 as an immunotherapeutic target, we next aimed to characterize the CD19 expression levels in CD19^+^ t(8;21) AML. To do so, we re-analyzed diagnostic flow cytometry data available for six CD19^+^ t(8;21) AML patients (patient #01-06) and one CD19^-^ t(8;21) AML patient (patient #07; control), which allowed for investigating the expression of CD19 on CD45^dim^SSC-A^low^CD34^+^ cells, and compared this to BMMC-derived CD19^+^ B- and CD19^-^ T cells (from patient #01) as a representative positive and negative control, respectively. The cell surface expression of CD19 in two patients (patients #01 and 02) approximated the expression level seen in CD19^+^ B cells, with a unimodal pattern (**Figure 3B**). In the remaining four CD19^+^ patients, we observed a continuum of CD19 expression levels on AML cells, ranging from the level seen in CD19^+^ B cells to that of T cells (range: 0 to 4 log; **Figure 3B**). Importantly, internal staining of the leukemic fusion protein RUNX1::ETO demonstrated a strong correlation with CD19 positivity (patient #01 and one additional primary BMMC sample: #08; **Figure 3C** and **S3A**). Given the success of blinatumomab in the treatment of BCP-ALL, we also investigated how the CD19 expression level on CD19^+^ t(8;21) AML samples (RL048 PDX and patient #08) compared to that on two primary BCP-ALL BMMC samples. While the CD19 MFI in AML patient #08 was lower compared to both BCP-ALL samples, the MFI of the AML PDX sample was just as high (ALL patient #02) or even higher compared to the BCP-ALL samples (patient #01) (**Figure 3D**).

**Figure 3.**
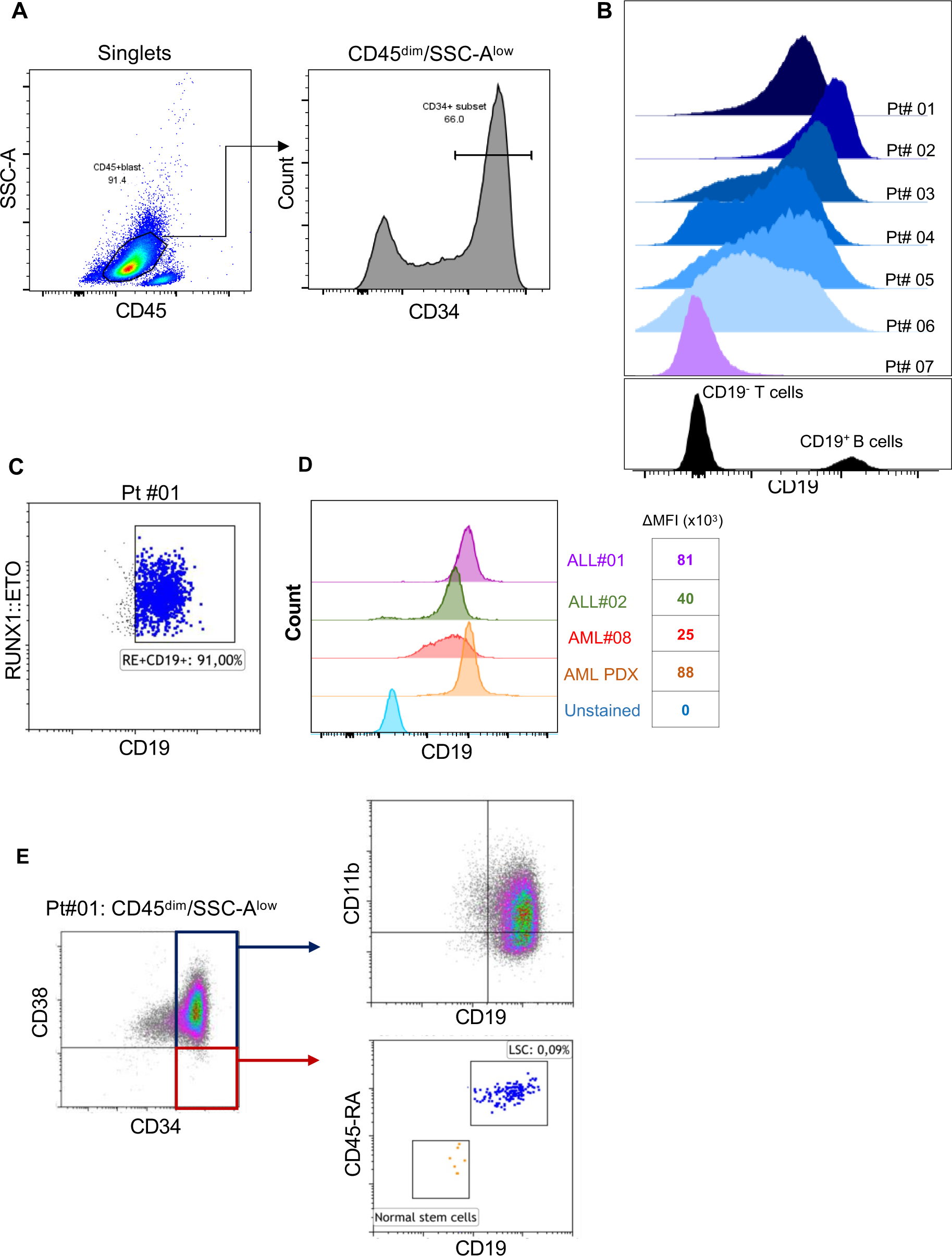
Overall and subpopulation-specific CD19 expression in CD19^+^ t(8;21) AML. (A) The gating strategy applied to myeloid blasts to identify CD19 positive populations. (B) Overview of the overall CD19 expression among CD45^dim^SSC-A^low^CD34^+^ blasts in the bone marrow. Data were retrieved from available diagnostic files of six CD19^+^ t(8;21) AML patients (patient #01-06) and one CD19^-^ t(8;21) AML patient (patient #07; reference) and were compared to the CD19 expression on T cells (as CD19^-^ control) and B cells (as CD19^+^ control). (C) Co-expression of CD19 and RUNX1::ETO among the myeloid blasts present in the bone marrow from patient #01. (D) Comparison of CD19 expression between AML PDX (RL048), one primary sample (patient #08), and two primary BCP-ALL samples. ΔMFI is calculated by subtracting the MFI of CD19 in stained samples from the corresponding unstained samples. (E) CD19 expression among leukemic stem cells (CD34^+^CD38^-^CD45RA^+^) and more mature subpopulations (CD34^+^CD38^+^CD11b^+^) phenotypes in patient #01. LSC: leukemic stem cell; PDX: patient-derived xenograft.

Since individual AML cells in the BM may vary in terms of maturation stages, targeting both immature and more mature AML cells is necessary for sustained therapeutic benefit.^38^ Therefore, we next sought to investigate the CD19 expression on different AML subpopulations, including CD34^+^CD38^-^ cells that encompass the leukemic stem cell (LSC) compartment, as well as those with a CD34^+^CD38^+^ phenotype. Performing flow cytometry on three samples (AML patients #01, #08, and RL048 PDX cells), we observed that nearly all CD34^+^CD38^-^ immature progenitors were positive for CD19 (**Figure 3E** and **S3BC**). These data are in line with previous data in two adults with t(8;21) AML showing that 77 and 91% of CD34^+^CD38^-^ cells were CD19^+^, respectively.^39, 40^ Furthermore, using a more extended flow cytometry panel for analysis of BMMCs from patient #01, we identified CD19 to be expressed on LSCs (CD34^+^CD38^-^CD45RA^+^) but not on normal stem cells (CD34^+^CD38^-^CD45RA^-^; **Figure 3D**). Similar to immature subpopulations, virtually all CD34^+^CD38^+^ blasts were positive for CD19. Moreover, we noted CD19 expression on both CD34^+^CD38^+^CD11b^+^ and CD34^+^CD38^+^CD11b^-^ cells, indicating CD19 expression on both more and less mature AML cells (**Figure 3D** and **S3B, S3C**). Taken together, these observations highlight that, in case of high overall AML CD19 expression, both primitive and more differentiated AML cells are potential targets of CD19-directed immunotherapies in CD19^+^ t(8;21) AML, encouraging exploration of their *ex vivo* killing efficiency.

### Blinatumomab is capable of activating T cells when bound to AML cells

Given the possibility of defective immune synapse formation between AML- and T cells, impairing proper T cell activation,^41^ we next investigated whether blinatumomab-mediated AML-T cell contact could facilitate the activation of T cells. Using genetically engineered Jurkat cells (CD3^+^CD4^+^) that express luciferase upon induction of the IL2 promoter following CD3 activation **(Figure 4A)**, we observed a dose-dependent increase in the luminescent signal in a co-culture of CD19^+^ AML PDX- and Jurkat cells **(Figure 4B)**, suggesting that blinatumomab can activate CD3 signaling in T cells by binding to CD19^+^ AML cells. To further validate this finding, we co-cultured healthy donor T cells with CD19^+^ AML PDX (RL048) cells in the presence or absence of blinatumomab for 48 hours. Addition of blinatumomab to the RL048 and T cell co-culture led to significant upregulation of the T cell activation markers CD25 (50% marker positivity) and CD137 (90% marker positivity), indicative of potent T cell activation (**Figure 4C**). In summary, these findings demonstrate that blinatumomab can activate T cells when bound to CD19^+^ AML cells. Based on the T cell activation assay, we identified 1 nM as the optimal concentration of blinatumomab to activate T cells in our co-cultures. Therefore, this concentration was used in subsequent *ex vivo* co-cultures involving blinatumomab.

**Figure 4.**
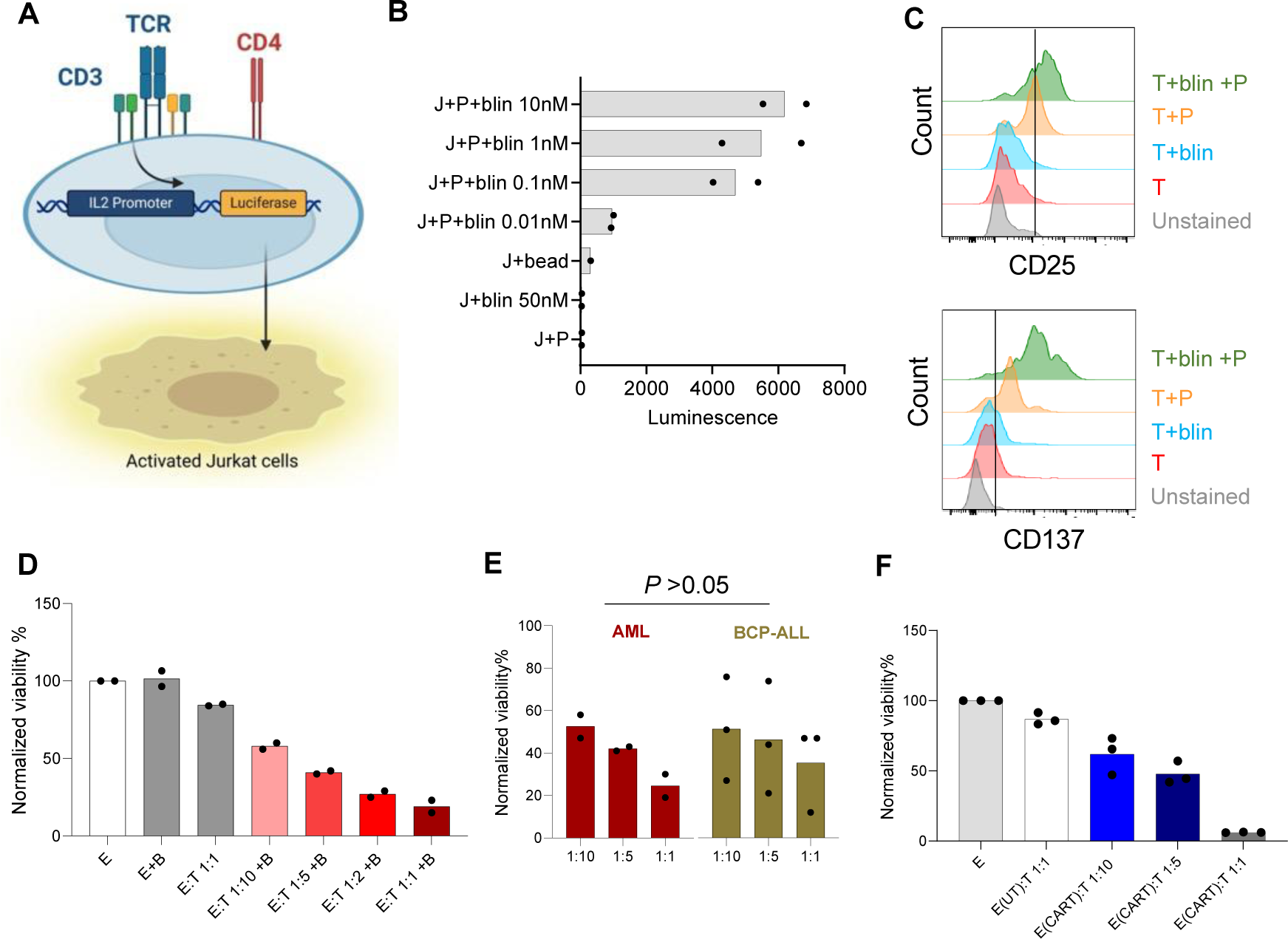
T cell activation and/or AML cell cytotoxicity mediated by blinatumomab and CAR T cells. (A) Illustration of the T cell activation bioassay. (B) The luminescent signal intensity upon addition of blinatumomab to CD19^+^ AML PDX and Jurkat cells (n=2 technical replicates). (C) Expression of the T cell activation markers upon addition of blinatumomab and/or PDX cells compared to healthy donor T cells alone. (D) Effect of 1 nM blinatumomab on the viability of PDX cells at various effector-to-target (E:T) ratios using healthy donor T cells after 48 hours. Data points represent technical replicates. (E) Comparison of blinatumomab-induced cytotoxicity in AML (n=2: patient #08 and PDX) and BCP-ALL patient samples (n=3), after 48 hours. Data represent mean ±SD. *t*-test was performed between each E:T ratio in AML versus BCP-ALL. (F) The viability of primary AML cells (patient #08) after 48 hours of co-culture with CAR T cells or untransduced T cells (control) at different E:T ratios. Data points represent technical replicates. J: Jurkat cells; P: PDX (patient-derived xenograft) cells; bead: CD3/CD28 Dynabeads; B or blin: Blinatumomab. E: effector (T cells); T: target (AML).

### CD19^+^ AML is sensitive to immunotherapy-mediated T cell cytotoxicity *ex vivo*

To determine whether AML cells were sensitive to blinatumomab-mediated T cell cytotoxicity, we co-cultured CD19^+^ AML PDX cells with or without healthy donor T cells for 48 hours, in the absence or presence of blinatumomab. Treatment of PDX cells with 1 nM of blinatumomab induced 40% AML cell killing at a low E:T ratio of 1:10 and almost 90% killing at an E:T ratio of 1:1 (**Figure 4D**). Importantly, absence of allogeneic T cells or blinatumomab led to no or negligible background killing (**Figure 4D**). We next compared the blinatumomab-mediated T cell killing efficiency between AML and BCP-ALL cells, at increasing E:T ratios of healthy donor T cells. Intriguingly, although we observed substantial variation in the killing efficiency among the three BCP-ALL samples, the observed AML cell killing was comparable to that in BCP-ALL in each E:T ratio (*P* > 0.05; **Figure 4E**).

In addition to blinatumomab, CD19-directed CAR T cells have been approved for the treatment of both pediatric and adult BCP-ALL.^13, 15^ Therefore, we assessed whether CD19^+^ t(8;21) AML cells were sensitive to CAR T cell-mediated cytotoxicity. Similar to blinatumomab, co-culture of CAR T cells with primary AML cells at a low E:T ratio of 1:10, led to 40% killing of AML cells within 48 hours, while AML cells were nearly completely eradicated at an E:T ratio of 1:1 (**Figure 4F**). Notably, AML cell viability remained constant in co-cultures with untransduced T cells from the same donor, indicating negligible background killing (**Figure 4F**). Taken together, these findings demonstrate that CD19-directed immunotherapies induce efficient killing of CD19^+^ AML cells *ex vivo*. These promising data prompted us to investigate whether the BM microenvironment of CD19^+^ t(8;21) AML patients is supportive of CD19-directed anti-tumor immunity.

### The composition of the bone marrow immune microenvironment of t(8;21) AML is comparable with non-leukemic controls but distinct from BCP-ALL

Previous work in AML, BCP-ALL, and various other cancers has shown that the efficacy of bispecific T cell-engagers and adoptive cell therapy largely depends on the presence and function of various immune cell populations in the tumor microenvironment.^8, 42–48^ To understand whether pediatric CD19^+^ t(8;21) AML patients may represent a subgroup with potential to respond to CD19-directed immunotherapies, we characterized their tumor immune microenvironment using immunogenomic computational approaches applied to diagnostic BM bulk RNA-seq data (**Figure 5A**). Towards this end, we deconvoluted the immune cell abundance in the BM of treatment-naïve CD19^+^ t(8;21) AML (n=5), CD19^-^ t(8;21) AML (n=5), other AML genotypes (n=30), and non-leukemic controls (n=4) using CIBERSORTx and the TIDE algorithm.^27, 30^ In an exploratory analysis, we did not detect differences in the estimated abundance of lymphoid subsets between CD19^+^- and CD19^-^ t(8;21) AML (**Figure S4A-G**). Likewise, no difference was observed among myeloid and stromal cell types (**Figure S4H-J**). Therefore, we considered these cases in aggregate for subsequent comparisons (referred to as t(8;21) group; n=10). We did not detect any differences in the abundance of microenvironmental subsets between both AML groups and non-leukemic controls (**Figure 5B-E, S4K-P**). In line with this, multiple RNA-based metrics related to immune function and -escape were similar among the three groups (**Figure 5F-I**). Indeed, T-and NK cell-related cytolytic activity (comprised of *GZMA*, *GZMH*, *GZMM, GNLY*, *PRF1*) (**Figure 5F**),^31^ a 172-gene immune effector dysfunction score (IED172) reflecting T- and NK cell exhaustion and senescence (**Figure 5H**),^32^ and HLA I and -II expression in AML patients did not differ from non-leukemic controls (**Figure 5H-I**).^31^ These data indicate that the BM immune microenvironment of t(8;21) AML at diagnosis does not harbor a particularly dysfunctional immune effector fraction nor is it highly immunosuppressive in comparison to non-leukemic controls, suggestive of low immune pressure in this AML subtype.

**Figure 5.**
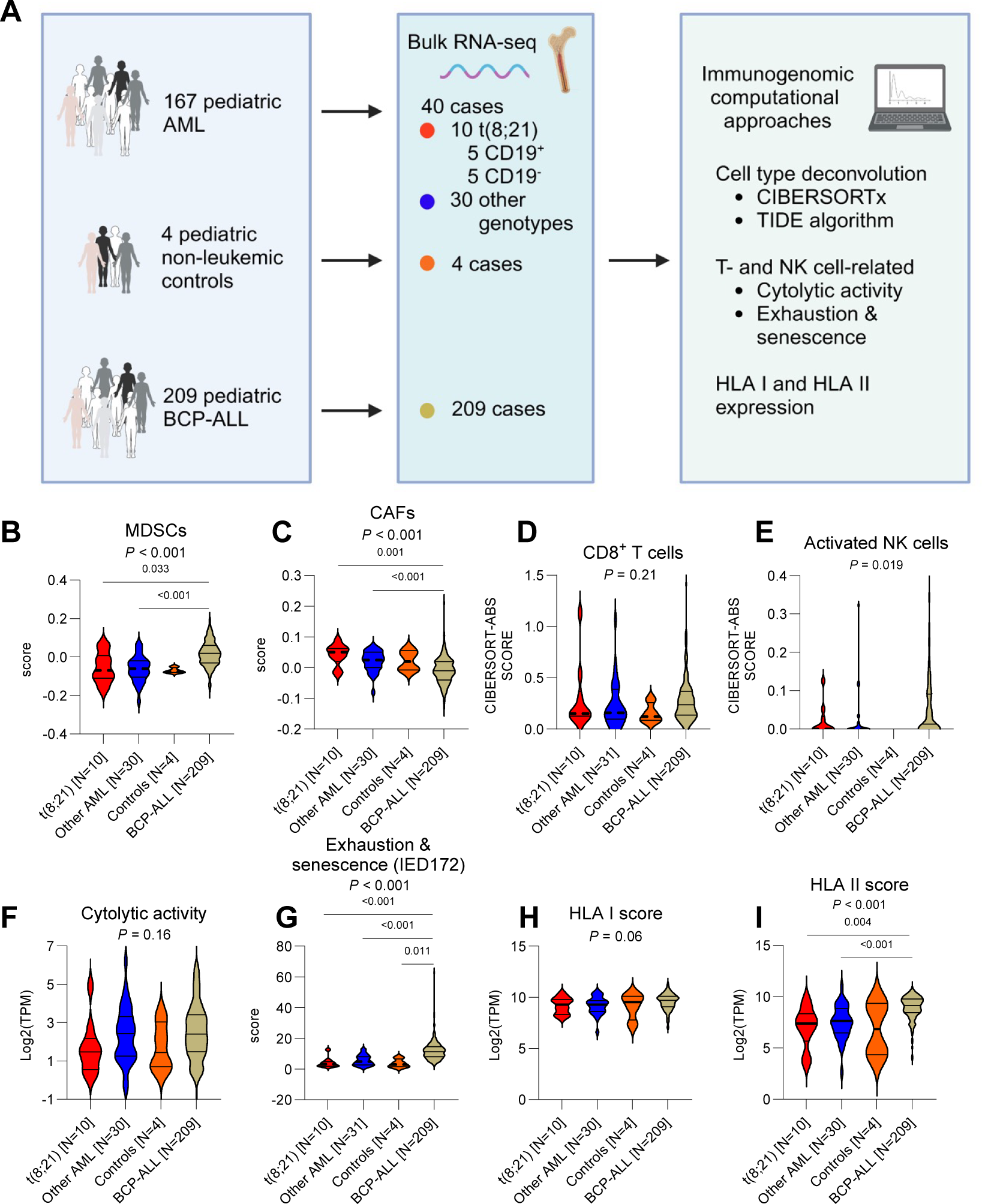
Characterization of the bone marrow immune microenvironment of t(8;21) AML using immunogenomic analyses. (A) Cohort overview for the characterization of the bone marrow (BM) immune microenvironment in pediatric AML, pediatric BCP-ALL, and non-leukemic controls. The non-leukemic controls are four pediatric patients with early-stage rhabdomyosarcoma without malignant BM infiltration (methods). (B-I) Comparison of the abundance of various cell populations and gene signatures among t(8;21) AML patients, AML patients with other cytogenetic alterations, BCP-ALL patients, and non-leukemic controls. Data are presented as median with quartiles and range. The statistical tests used include the Kruskal-Wallis test with Dunn’s post-hoc test for multiple comparisons. In case two *P* values are shown, the upper one indicates the result of the Kruskal-Wallis test, while the lower *P* values indicate the result of Dunn’s test. MDSC: myeloid-derived suppressor cell; CAF: cancer-associated fibroblast; NK: natural killer; IED172: 172-gene immune effector dysfunction score; HLA: human leukocyte antigen.

As CD19-directed immunotherapies have led to impressive and durable responses in pediatric BCP-ALL, we next applied our immunogenomic approach to investigate how the diagnostic BM immune microenvironment of pediatric t(8;21) AML (n=10) compared to that of pediatric BCP-ALL (n=209; **Figure 5A**). As anticipated because of the B cell origin of BCP-ALL, we detected a significant enrichment in naïve B cells in comparison to t(8;21) AML (**Figure S4K**). Furthermore, BCP-ALL cases had a significantly higher abundance of MDSCs and were enriched for T- and NK cell exhaustion and senescence, potentially reflecting a prior T- and NK cell response rendered dysfunctional (**Figure 5B and G**). On the other hand, CAFs, memory B cells, and plasma cells were significantly increased in t(8;21) AML compared to BCP-ALL, albeit absolute differences were minimal for the latter two (**Figure 5C** and **S4L-M**). Whereas no differences in HLA I expression were detected, HLA II expression was significantly increased in BCP-ALL compared to t(8;21) AML (**Figure 5H-I**), which is likely related to the antigen presenting cell-origin of BCP-ALL cells.^31^ Altogether, our immunogenomic approach revealed that the BM microenvironment in pediatric t(8;21) AML is comparable to that of non-leukemic controls but, at least in part, distinct from that of pediatric BCP-ALL.

### Autologous T cells from t(8;21) AML patients are functional and induce cytotoxicity upon activation by blinatumomab

Following the characterization of the BM immune microenvironment in t(8;21) AML patients, we sought to evaluate the efficacy of blinatumomab-mediated killing of CD19^+^ t(8;21) AML cells by autologous T cells present within BMMC samples (n=2), and to compare this to primary BCP-ALL (n=3). Such an autologous killing assay would reveal the functionality of AML T cells compared to those present in BCP-ALL, at the naturally occurring E:T ratio and in the presence of other BMMCs, approximating the *in vivo* composition. The two AML samples contained 4% and 8% CD3^+^ T cells, respectively, while all three BCP-ALL samples harbored nearly 3% CD3^+^ T cells. Interestingly, addition of blinatumomab (1 nM) to 1 x 10^5^ BMMCs led, in 48 hours, to a reduction in AML cell viability of approximately 50% compared to the viability in the absence of blinatumomab, which was comparable to that seen in the BCP-ALL samples (*P* > 0.05; **Figure 6A**). To confirm that the reduced AML cell numbers were due to blinatumomab-mediated T cell killing, we measured IFN-γ secretion and the expression of activation markers on the autologous T cells. In both AML samples, we found that blinatumomab induced a significant increase in IFN-γ secretion, with the extent proportional to the abundance of T cells in the BM (**Figure 6B**). In addition, the expression of the T cell activation markers CD25 and CD137 on BM T cells increased substantially in response to blinatumomab (**Figure 6C**). For patient #01, matched PB was also available, which allowed for a co-culture of PB-derived T cells and autologous BMMCs for 48 hours with or without blinatumomab. Consistent with the findings with autologous BM T cells, autologous PB T cells, co-cultured with matched BMMCs, showed substantial activation upon treatment with blinatumomab (**Figure 6D**). Accordingly, PB-derived T cells demonstrated effective AML cell killing (**Figure 6E**). Furthermore, this was accompanied by substantial IFN-γ secretion (**Figure 6F**) and increased T cell numbers after treatment (**Figure 6G**). Overall, these findings demonstrate that autologous T cells from AML patients are capable to induce cytotoxicity upon binding to T cell-engagers, encouraging the exploitation of CD19 as an immunotherapy target in pediatric CD19^+^ t(8;21) AML.

**Figure 6.**
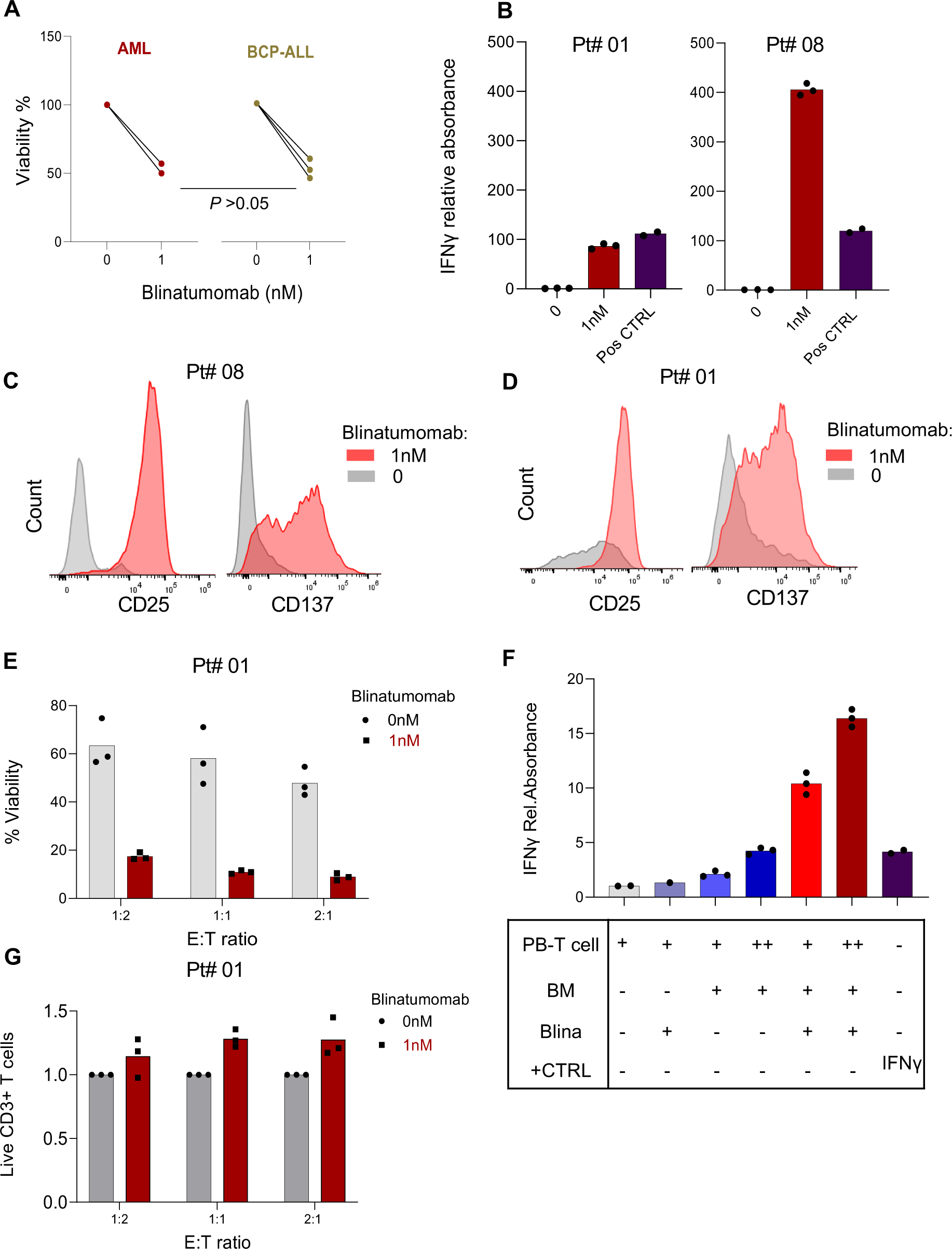
T cells from bone marrow and peripheral blood of t(8;21) patients are functional and actively induce cytotoxicity upon blinatumomab administration. (A) Cytotoxicity of autologous bone marrow (BM)-derived T cells upon addition of 1 nM blinatumomab to BM mononuclear cells (MCs) from AML (n=2) and BCP-ALL (n=3) samples after 48 hours. t-test was performed to compare results for blinatumomab-treated AML and BCP-ALL samples (B) Relative interferon (IFN)-γ measurement in the supernatant of two BMMC AML samples; positive control: 125 pg/mL recombinant IFN-γ protein, n= 3 technical replicates. (C-D) Changes of the activation markers on CD3^+^ T cells present in the BMMC sample (patient #08) (C) or derived from peripheral blood (patient #01) (D). (E) Cytotoxicity of autologous peripheral blood-derived CD3^+^ T cells upon co-culture with matched BMMCs from patient #01 after 48 hours; n=3 technical replicates. (F) Relative IFN-γ measurement in the supernatant of BMMCs from patient #01 upon co-culture with matched BMMC cells at different E:T ratios. The + and ++ in the table beneath show the ratio of T cells to AML cells, respectively. Absorbance values were normalized to the corresponding value with CD3^+^ T cells alone; positive control: 125 pg/ml recombinant IFN-γ, n= 3 technical replicates. CTRL: control. (G) Changes in the number of peripheral blood-derived CD3^+^ T cell numbers in the presence or absence of 1 nM blinatumomab and in co-culture with BMMCs from patient #01 after 48 hours. Cell numbers for each condition (varying E:T ratios) were normalized to the corresponding condition without blinatumomab.

## Discussion

Repurposing immunotherapies that have been approved for other hematological malignancies may not only accelerate the realization of potential clinical benefits, it can also reduce the inherent risks and delays associated with introducing novel agents. The success of CD19-directed immunotherapies in BCP-ALL, as well as in two adults with relapsed CD19^+^ t(8;21) AML,^17, 18^ prompted us to investigate whether CD19 could be a valuable immunotherapy target in pediatric AML. Our study reveals that a subset of pediatric AML patients, in particular those with t(8;21) AML, express CD19 at diagnosis, consistent with findings in adult AML.^33^ Importantly, the extent of CD19 expression on AML cells among those classified as having CD19^+^ AML was heterogeneous, indicating that not all CD19^+^ t(8;21) AML patients may be suitable candidates for CD19-directed immunotherapy. In those with unimodal and high CD19 expression, CD19 was expressed on nearly all CD34^+^CD38^-^ and CD34^+^CD38^+^ subpopulations, suggesting potential elimination of these AML subpopulations through CD19-directed immunotherapies. Our *ex vivo* studies revealed that blinatumomab was able to induce AML cell killing with an efficacy comparable to that seen in BCP-ALL. Moreover, CAR T cells could effectively eliminate CD19^+^ AML cells *ex vivo*. Lastly, T- and NK cells in the bone marrow of pediatric t(8;21) AML appeared to be less exhausted and senescent in comparison to pediatric BCP-ALL. Collectively, our study demonstrates the potential of CD19-directed immunotherapies for the treatment of pediatric CD19^+^ AML.

While pediatric t(8;21) AML has a favorable prognosis following current chemotherapy regimens (5-year OS rate of nearly 90%),^49^ alternative therapies are needed to reduce treatment-related toxicity in newly diagnosed patients, and to improve outcomes in relapsed and refractory disease. The comparable *ex vivo* blinatumomab-mediated killing efficiency in CD19^+^ t(8;21) AML and BCP-ALL suggests that the successes observed with CD19-directed immunotherapies for BCP-ALL may be seen in CD19^+^ t(8;21) AML as well. Given that immunotherapies work best at a favorable E:T ratio, a potential setting for the use of CD19-directed immunotherapies could be that of minimal residual disease (MRD)-positivity before allo-SCT or other cellular therapies with curative intent.^50^ Furthermore, we envision that these therapies could serve as an alternative to intensive chemotherapy in case of excess toxicity, or as a life-prolonging treatment when curative options are no longer viable. Of relevance, given the heterogeneous expression of CD19 in those classified as having CD19^+^ AML, flow cytometry should be used to assess the fraction and intensity of AML cells positive for CD19. Moreover, data from our study and others show that a subset of patients with CD19^-^ AML at diagnosis gained CD19-expression at relapse, highlighting another subgroup that could potentially benefit from CD19-directed immunotherapies as well.^51^

In addition to our *ex vivo* studies, our characterization of the BM immune microenvironment provides insight into the *in vivo* setting, which may further contribute to identifying patients that are likely to benefit from these immunotherapies. Interestingly, our immunogenomic analyses revealed that the BM immune microenvironment in pediatric t(8;21) AML was highly similar to that of non-leukemic controls, suggestive of low immune pressure. In addition, pediatric t(8;21) AML appeared to have a less exhausted and senescent T- and NK cell compartment in comparison to pediatric BCP-ALL. As T- and NK cell exhaustion and senescence have recently been linked to resistance to bispecific antibodies and immune checkpoint inhibitors, the more inert T- and NK cell state in t(8;21) AML could be a favorable starting point for response to CD19-directed immunotherapies.^32^

A limitation of our study is the relatively small number of CD19^+^ AML samples available for our *ex vivo* studies. Nonetheless, the observed efficacy of CD19-directed immunotherapies was highly similar among the investigated samples, indicating robustness of our findings.

In conclusion, the high frequency of CD19 expression in pediatric t(8;21) AML, in combination with our *ex vivo*- and immunogenomic studies, suggests that CD19 can be exploited as an immunotherapy target in t(8;21) pediatric AML, and potentially in other AML subtypes exhibiting CD19 positivity as well. The eagerly anticipated results of three clinical trials that are investigating CD19-directed immunotherapies in R/R adult (NCT04257175 and NCT03896854) and pediatric AML (NCT02790515) will shed further light on the potential of these therapies in AML. In addition, we have initiated an international registry for pediatric AML patients treated with CD19-directed immunotherapies, which will simultaneously generate relevant knowledge regarding the efficacy and safety of these therapies in the pediatric population.

## Acknowledgements

We would like to express our gratitude to all patients and their parents/caregivers for their participation in the research. We are grateful to Anja de Jong (Princess Máxima Center) for acquiring flow cytometry data; Zinia Kwidama for providing AML samples (UMC Amsterdam, location VUmc, The Netherlands), and Tom Reuvekamp and Marisa Westers for fruitful discussions of the data. Feeders for rapid T-cell expansion were kindly provided by dr. Zsolt Sebestyen (UMC Utrecht, The Netherlands). Figures were created with BioRender.com.

## Author contributions

F.B., N.W., G.K., and O.H. formulated the study concept and designed experiments. The experiments were performed by F.B., J.B.K., N.W., T.M., M.A., and E.D. CAR T cell generation was performed by N.D., and A.M.C. The AML medium was optimized by A.K.H. Data interpretation was performed by F.B., N.W., and J.B.K. Co-supervising the panel design for identification of leukemic cells in killing assays of primary AML samples was done by J.C. CAR T cell production was supervised by S.N. and J.K. The manuscript was written by F.B, J.B.K., N.W., and O.H. together with T.M. The study was supervised by K.K., C.Z., G.K., and O.H. All authors read and approved the final version of the manuscript.

## Data availability

Sequencing data can be accessed from the Gene Expression Omnibus (GSEXXX; normalized counts [GSE IDs will be available upon publication]. Raw sequencing data requests should be addressed to and will be fulfilled by the corresponding author.

