## Supplemental Data and Figures for "Repurposing CD19-directed immunotherapies for pediatric t(8;21) acute myeloid leukemia"

to

#### MATERIALS AND METHODS

##### Ethical regulations

All patients provided written consent for banking and use of clinical data, BM- and peripheral blood (PB) mononuclear cells (MCs), and bulk RNA-sequencing (bulk RNA-seq) data for research purposes.

##### AML treatment regimen

All AML patients were treated according to the NOPHO-DBH AML-2012 protocol[1]. Protocol treatment consisted of two induction courses (MEC: mitoxantrone-etoposide-cytarabine and ADE: cytarabine-daunorubicin-etoposide) and three consolidation courses (HAM: high-dose cytarabine-mitoxantrone, HA3E: high-dose cytarabine-etoposide, and FLA: fludarabine-cytarabine). Initially, induction course 1 consisted of either MEC or DEC (daunorubicin-etoposide-cytarabine) and induction course 2 of ADE or FLAD (fludarabine-cytarabine-daunorubicin) randomizations. However, MEC and ADE outperformed the other courses and therefore became the standard treatment as of January 2019.[1] If possible, high-risk patients (those with suboptimal response to standard therapy or those with a *FLT3*-ITD without a concurrent *NPM1* mutation) received an allogeneic stem cell transplantation (allo-SCT) after HAM. In addition, standard risk patients with an inv(16) received two instead of three consolidation courses (omission of HAM).

##### Definitions and response criteria according to NOPHO-DBH AML-2012

- Complete remission is defined as <5% leukemic cells in the BM, as determined by either morphology or flow cytometry, with clear evidence of hematological regeneration. This includes levels of ANC ≥0.5 x 10^9/L, platelets ≥50 x 10^9/L, and no signs of leukemia elsewhere.

- Refractory disease (RD) was defined as the presence of ≥ 5% leukemic cells in the BM after the second induction course.
- The term relapse was reserved for patients that relapsed, according to the following definitions, after they achieved CR. BM relapse was defined as the presence of ≥5% leukemic cells in the BM. Molecular relapse was defined as a more than tenfold increase (> one-log) in transcript level. CNS relapse was defined as the presence of leukemic cells in the CNS detected on cytospin preparations of CSF with ≥ 5 white blood cells/μL or the presence of a tumor mass in the CNS with identical or very similar characteristics compared to the original leukemic population. Extramedullary relapse was defined as the presence of a tumor mass outside of the BM with identical or very similar characteristics compared to the original leukemic population.
- Early death was defined as any death occurring in patients before achieving remission unless they have been classified as having resistant disease.
- Event-free survival (EFS) was defined as the time from diagnosis until the first event, or the date of last follow-up.
- Overall survival (OS) was defined as the time from diagnosis until death, or the date of last follow-up.

**Flow cytometry data: CD19 positivity**

Using collected reports of flow cytometry data, the CD19 positivity was determined as follows: leukemic cells were considered $CD19^+$ if the difference in median fluorescence intensity (MFI) between the leukemic blasts and the healthy population was greater than 3-fold. Accordingly, a difference below 3-fold was considered negative. Importantly, in cases were CD19 expression was multimodal, the MFI of the positive part was considered.

**CD19 expression analysis**

To determine the CD19 expression on myeloid blasts, the following antibodies were used (all antibodies were purchased from BioLegend unless specified otherwise): CD19-BV421 or PE (SJ25C1), CD45-FITC (HI30), CD34-BV605 (BD-Biosciences, 8G12), CD38-PerCP-Cy5.5 (HIT2), CD45RA-BV785 (HI100), CD11b-PE (ICRF44), CD33-APCFire750 or CD33-BV421 (P67.6), and CD20-FITC (2H7; used for exclusion of $CD19^+$ B cells). Expression of CD19 was compared to T- and B cells as $CD19^-$ and $CD19^+$ controls, respectively.

**Co-expression of CD19 with RUNX1::ETO**

Primary AML cells (2 x $10^5$) were stained with CD19_BV421(1:50) and live/dead marker viakrome 808 (1:50) for 30 min at room temperature (RT) and were washed with PBA-F (PBS with 0.025% BSA, 0.02% NaN3, 1% FBS) twice and pelleted in v-bottom plates. Cells were resuspended in 100 µL fixation buffer (BD Cytofix, BD Biosciences) by vortexing and incubated for 15 min at 4 °C. Cells were washed twice, transferred into 1.5ml microtubes, and permeabilized by slowly adding 500 µL ice cold (kept at -20 °C) BD Phosflow™ Perm Buffer III (BD Biosciences) while vortexing for 30 min on ice. After washing with PBA-F, blocking solution containing 10% donkey serum in PBAF was used for 1 hr at RT. Cells were then resuspended in anti-ETO polyclonal antibody (1:250 in PBA-F, Invitrogen, Cat# PAS-79943) for 1 hr at RT. After 2 washing steps, cells were stained with secondary antibody (donkey anti-rabbit IgG labelled to AF555, Invitrogen, Cat# 406412) in PBA-F (1:2000) for 45 min in the dark. Cells were washed twice and resuspended in PBA-F before performing flow cytometry using the Cytoflex LX (Beckman Coulter).

##### *Ex vivo* primary AML and T cell cultures

CD19$^+$ primary BMMCs from t(8;21) AML patients and RL048 PDX cells were resuspended at 0.5-1.0 x $10^6$ cells/mL in an in-house optimized AML medium containing SFEMII (STEMCELL Technologies) supplemented with 10 ng/mL FLT3-L, 10 ng/mL granulocyte-macrophage colony-stimulating factor (GM-CSF), 10 ng/mL IL-3, 100 ng/mL TPO, 150 ng/mL SCF (all from PeproTech), 750 nM SR1 (BioGems), 1.35 µM UM729 (STEMCELL Technologies). T cells (number of cells was dependent on the E:T ratio) were resuspended in RPMI plus GlutamaxTM (Gibco), 5% heat-inactivated human serum (Sigma) + 50 IU IL2, 5 ng/mL IL7, and 5 ng/mL IL15 (all from Peprotech; for the CAR T cells, IL15 was purchased from Miltenyi Biotec). For killing assays, a 1:1 mixture of AML and T cell medium (2x concentrated) was used.

##### *Ex vivo* T cell killing assays

The leukemic fraction within AML BMMC samples was defined as CD45$^{dim}$SSC-A$^{low}$CD19$^+$CD33$^+$CD34$^+$CD3$^-$CD20$^-$ cells, while for BCP-ALL BMMC samples this was defined as CD19$^+$CD10$^+$CD22$^+$CD3$^-$ cells. To distinguish remaining live RL048 PDX cells from co-cultured T cells after the killing assay, T cells were pre-stained with 1 µM CellTrace Violet (CTV, Invitrogen) according to manufacturer's instructions, before adding them to the PDX cells. By the end of the experiment, Sytox red (dilution 1:2000) was added to exclude dead cells, and the percentage of live CTV$^-$ cells was used to determine the viability of RL048 PDX cells. For allogeneic killing assays with BMMCs, live myeloid blasts were defined as Viakrome808$^-$CD45$^{dim}$CD33$^+$CD34$^+$CD3$^-$CD20$^-$, while live lymphoid blasts were defined as Viakrome808$^-$CD10$^+$CD22$^+$CD3$^-$. To calculate background killing, sample viability in experiment conditions that included blinatumomab was normalized to the viability in the absence of blinatumomab. In allogeneic killing assays with CD19-directed CAR T cells, the same approach was used and included a co-culture with untransduced

T cells from the same donor to determine the background killing. Further details on the generation of CAR T cells can be found below.

**T cell activation assay**

T Cell Activation Bioassay (IL2 promoter kit, Cat# J1651, Promega) was used to assess the ability of AML cells to activate CD3 signaling in the presence of blinatumomab. The kit consists of a Jurkat lymphoma cell line ($CD3^+CD4^+$) engineered to express luciferase upon IL2-promoter activation. For the T cell activation assay, Jurkat and RL048 PDX cells were resuspended at $4*10^6$ cells/mL. As a positive control, CD3/CD28 Dynabeads (Gibco™) were used to activate CD3 signaling in Jurkat cells in the absence of blinatumomab or AML cells. From the suspension of each cell type, 25 µL was added to each well of a flat-bottom 96-well plate. Subsequently, 25 µL of RPMI or blinatumomab concentrations (3x concentrated) were added to each well. Cells were incubated for 6 hours at 37°C. Then, the luciferase assay and substrate were mixed, aliquots were made in amber color-tubes, and maintained at -20°C. Two hours before the assay measurement, aliquots were moved to RT. After 6 hours, the plate was removed from the incubator and equilibrated at RT for 10-15 minutes. 75 µL luciferase was added to each well and incubated for 5 minutes. Next, the plate was centrifuged for 3 minutes at 1000g, and 100 µL of supernatant was added to a white-opaque flat-bottom 96-well plates. Luminescence was measured within 10 minutes of luciferase addition, with an integration time of 500 ms. The concentration of blinatumomab to activate T cells in subsequent experiments was determined using this T cell activation assay. For assessing the activation of healthy donor T cells upon co-culture with $CD19^+$ AML and blinatumomab, flow cytometry was used. Following debri/doublet/dead cell removal, CD25 and CD137 positivity was assessed on a population positive for $CD3^+$ and $CD4^+$ or $CD8^+$.

**Interferon-γ secretion analysis**

Supernatants of AML-T cell co-cultures were collected at 48 hours and used to measure the interferon (IFN)-γ secretion using enzyme-linked immunosorbent assay (ELISA) with the Human IFN-γ uncoated ELISA kit (Invitrogen, 88-7316), according to manufacturer's instructions.

**Generation of CD19-directed CAR T cells**

The CAR19 gene was constructed by linking the FMC-63 single chain variable fragment (GenBank ID: ADM64594.1) to a CD8 hinge (AA 138-206, Ref sequence ID NP_001759.3) and transmembrane domain, 4-1BB (CD137, AA 214-241 UniProt sequence ID Q07011) transactivation domain, and CD3zeta signaling domain (CD247, AA 52-121, Ref sequence ID: NP_000725.1). The sequence was codon optimized and synthesized (Genscript), after which it was cloned in to the pCCL-cPPT-hPGK-ORF-bPRE4-SIN lentiviral transfer vector.[2] Lentiviral particles were produced using standard calcium phosphate transfection (Clontech, 631312) of HEK-293T-cells with the pMDL-g/pRRE, pMD2-VSVg, and pRSV-Rev plasmids.[3, 4] Transducing units per mL of concentrated vector batches were determined using serial dilution on Jurkat cells (ATCC, TIB-152) followed by flow cytometric analysis using a biotin-conjugated CD19 CAR detection reagent (Miltenyi Biotec, 130-129-550) with APC-conjugated streptavidin (Biolegend, 405207). Findings were confirmed using a serial dilution on HeLa cells followed by a qPCR to quantify the vector copy number of the construct within the DNA. Peripheral blood mononuclear cells were separated from blood using Ficoll-Paque PLUS separation (Cytiva, 17-1440-03), after which T cells were isolated with CD3 microbeads according to manufacturer's instructions (Miltenyi Biotec 130-097-043). T cells were activated and transduced at a multiplicity of infection of 10 according to a previously established protocol.[2] Six days after transduction, cells were sorted for CAR T receptor expression using a panel with the biotin-conjugated CD19 CAR detection reagent, PE-conjugated anti-biotin (Miltenyi Biotec,130-110-951), and eFluor (Invitrogen, 65-

0864-14). Cells were sorted on a Sony SH800S Cell Sorter. After sorting, cells were expanded using a previously established rapid T-cell expansion protocol.[5] Untransduced CD3 T cells undergoing rapid expansion were used as controls. Before co-culture with leukemic cells, T cells were resuspended in RPMI plus Glutamax™ (Gibco), 5% heat-inactivated human serum (Sigma) $^+$ 50 IU IL2, 5 ng/mL IL7, and 5 ng/mL IL15 (all from Peprotech; for the CAR T cells, IL15 was purchased from Miltenyi Biotec). For the autologous killing assays, the CD3$^+$ T cells from patients' PBMCs were isolated with CD3 microbeads and resuspended in the above-mentioned T cell medium.

**Bulk RNA-sequencing**

Briefly, total RNA was isolated from viably frozen BMMCs (derived from BM aspirates), RNA-seq libraries were generated from 300 ng RNA, and sequenced with NovaSeq 6000 (2x150 bp; Illumina). After pre-processing, RNA-sequencing yielded 80-100 million raw reads. Reads were aligned to GRCh38[6] and gencode version 31.[7]

To compare gene expression profiles of CD19$^+$ and CD19$^-$ t(8;21) AML, BM samples with a blast percentage ≥ 75% were selected out of all t(8;21) AML patients in the total cohort (n=4; 3 CD19$^+$ and 1 CD19$^-$ cases). In addition, to increase the cohort size for this analysis, we acquired publicly available BM bulk RNA-seq data from newly diagnosed AML patients (n=5 t(8;21) patients with BM blasts ≥ 75%; three CD19$^+$ and two CD19$^-$ cases; TARGET-AML cohort).[8] These five cases were selected based on the expression values (TPM) of *CD19* and typical B cell-related genes (*CD20* (also known as *MS4A1*) and *CD22*). CD19 positivity was determined as a more than three-fold higher expression of *CD19* compared to both *CD20* and *CD22*. CD19 negative cases were selected based on both low *CD19 and CD20/CD22* values (TPM below 1). Consequently, we identified the samples PARENB, PARGVC, and PASHBI to be CD19$^+$ t(8;21) samples, while PANDER and PATDHA were regarded as CD19$^-$ t(8;21) samples. Differential gene expression analysis including batch correction according to Combatseq was performed using DEseq2 in R (version 4.3.1) with a false-discovery rate (FDR) cutoff of 0.05 and minimum fold change of 2.[9, 10] Gene set enrichment analysis (GSEA) was performed using the software developed by UC San Diego (San Diego, CA) and the Broad Institute (Boston, MA) with the KEGG, Hallmarks, and Reactome gene sets.[11-17]

To infer the abundance of lymphoid populations from bulk RNA-sequencing data, we harnessed CIBERSORTx and focused on cell subsets that were previously validated in AML by flow cytometry and/or single-cell RNA-sequencing (naïve B cells, memory B cells, plasma cells, follicular helper T cells, resting $CD4^+$ memory T cells, $CD8^+$ T cells, and activated NK cells).[18, 19] Furthermore, we utilized the TIDE algorithm to infer the abundance of several immunosuppressive myeloid and stromal cell types (M2-like macrophages compared to M1-like macrophages (termed M2-predominance), cancer associated fibroblasts (CAFs), and myeloid-derived suppressor cells (MDSCs)), as this approach was developed specifically for five cancer types including AML.[20] For CIBERSORTx, count data were normalized to transcripts per million (TPM) and for TIDE, several normalization measures were applied to TPM data according to developers' instructions. In addition, we used several RNA-based metrics related to immune function and -escape: the cytolytic score,[21] the immune effector dysfunction score,[22] and HLA I and -II scores.[21] All AML cases that had bulk RNA-seq data available were part of the total cohort of 167 patients. For BCP-ALL, bulk RNA-seq data of 209 treatment-naïve patients were acquired based on availability in the institution's biobank without any selection for gender, age, cytogenetics, mutational status, or outcomes. Non-leukemic controls were BM samples from pediatric patients with treatment-naïve, early stage rhabdomyosarcoma with no BM infiltration that were previously profiled at our institution as described above.[23]

**Statistical analyses**

Probabilities of event-free survival (pEFS) and overall survival (pOS) of the overall cohort were estimated using the Kaplan-Meier method. Differences between two independent groups were compared using the t-test or Mann-Whitney test, depending on the distribution of the data. Differences between three or more groups were compared using the ordinary one-way ANOVA test in combination with Tukey's multiple comparisons test for data with Gaussian distribution of residuals, or using the Kruskal-Wallis test in combination with Dunn's multiple comparisons test for data without such a distribution.

#### TABLES AND FIGURE LEGENDS

##### Supplementary Tables

**Supplementary Table 1: Overview of patient characteristics of bulk RNA-seq AML cohort.**

**Supplementary Table 2: Overview of patient characteristics of bulk RNA-seq BCP-ALL cohort.**

**Supplementary Table 3: Overview of differentially expressed genes between CD19$^{+}$- and CD19$^{-}$ t(8;21) AML.**

Note: all Supplementary Tables are provided in an excel file.

**Supplementary Figure 1. Impact of CD19 expression on patient outcomes in t(8;21) AML.**

(A-B) Probability of event-free survival (A) and overall survival (B) of CD19$^{+}$- and CD19$^{-}$ t(8;21) AML patients, following standard chemotherapy treatment (Supplementary Methods).

**Supplementary Figure 2. Altered pathway activities between CD19$^{+}$- and CD19$^{-}$ t(8;21) AML.**

(A-F) Gene set enrichment analysis of bulk RNA-seq data showing depleted gene sets and pathways in CD19$^{+}$- compared to CD19$^{-}$ t(8;21) AML.

**Supplementary Figure 3. CD19 expression on AML subpopulations.** (A) Co-expression of CD19 and RUNX1::ETO in BMMCs from patient #08. (B) Gating strategy and CD19 expression among CD34$^{+}$CD38$^{-}$, CD34$^{+}$CD38$^{+}$, and CD34$^{+}$CD38$^{+}$CD11b$^{+}$ cells in BMMCs from patient #08. (C) Gating strategy and CD19 expression among CD34$^{+}$CD38$^{-}$, CD34$^{+}$CD38$^{+}$ and CD11b$^{+}$ cells from RL048 patient-derived xenograft (PDX) cells.

**Supplementary Figure 4. Characterization of the bone marrow immune microenvironment of t(8;21) AML using immunogenomic analyses.** (A-J) Comparison of the deconvoluted abundance of various cell populations among CD19$^{+}$- and CD19$^{-}$ t(8;21) AML. Data are presented as mean with standard deviation. The Mann-Whitney test was used to test for statistical differences between groups. M2: anti-inflammatory macrophage subtype; CAF: cancer-associated fibroblast; MDSC: myeloid-derived suppressor cell. (K-P) Comparison of the deconvoluted abundance of various cell populations among t(8;21) AML patients, AML patients with other cytogenetic alterations, BCP-ALL patients, and non-leukemic controls. Data are presented as median with quartiles and range. The statistical tests used include the Kruskal-Wallis test with Dunn's post-hoc test for multiple comparisons. In case two *P* values are shown, the upper one indicates the result of the Kruskal-Wallis test, while the lower *P* values indicate the result of Dunn's test.

### Supplementary Figure 1

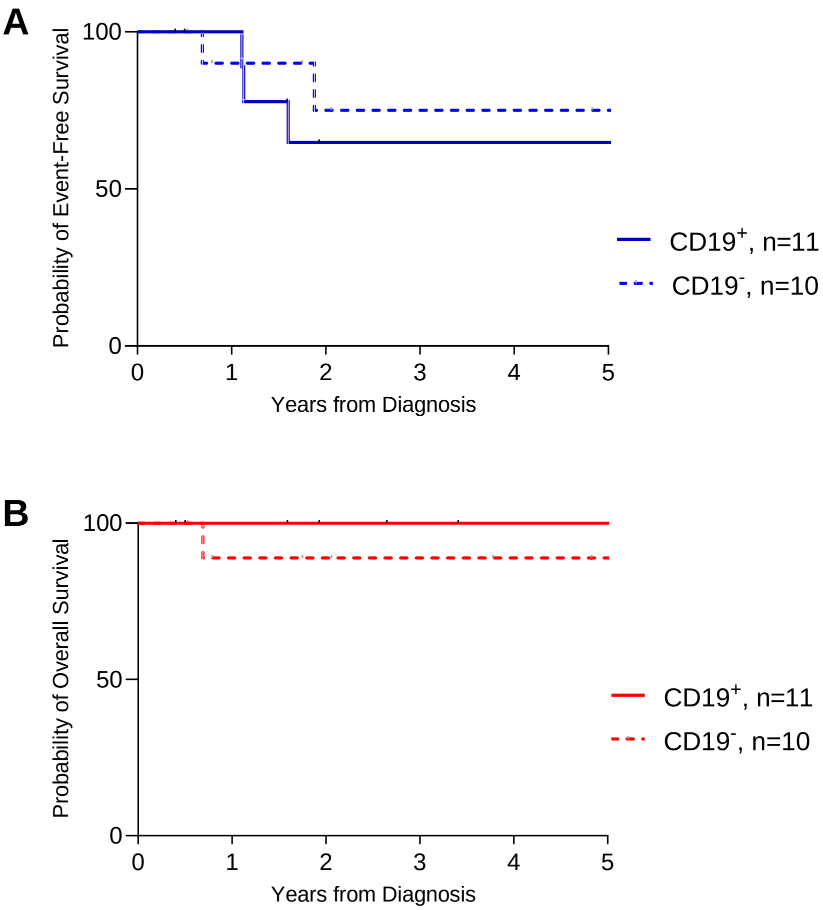

### Supplementary Figure 2

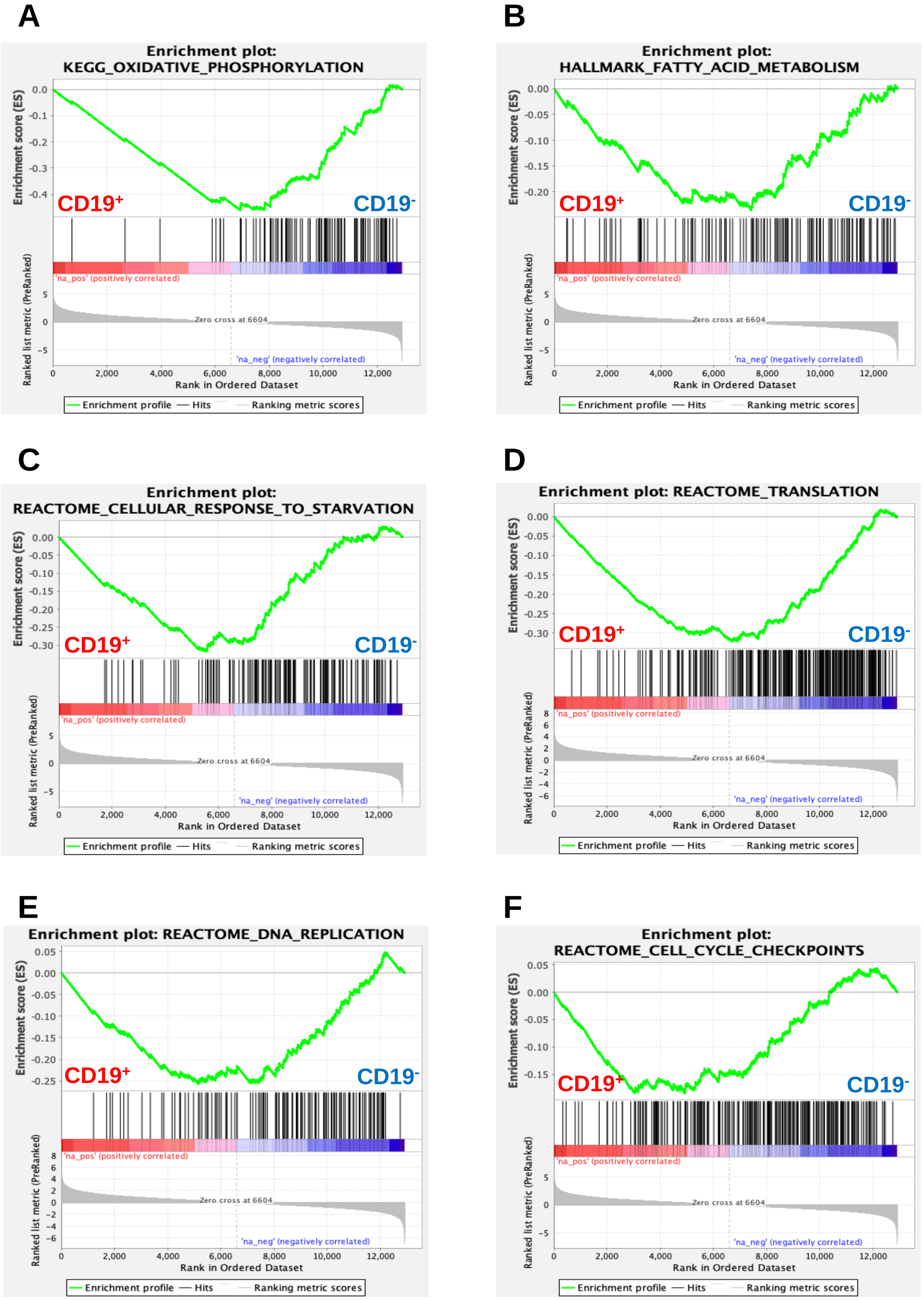

### Supplementary Figure 3

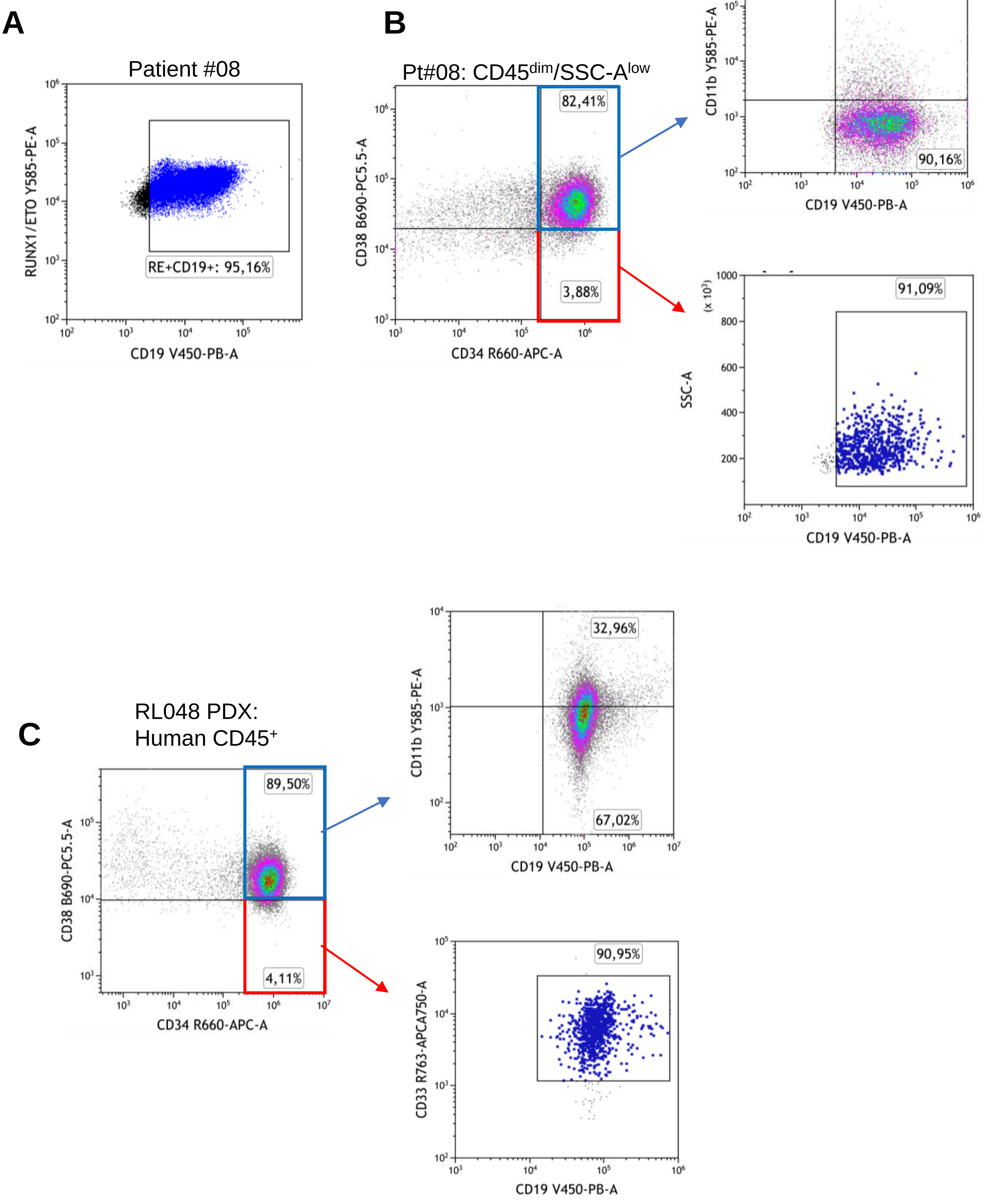

### Supplementary Figure 4

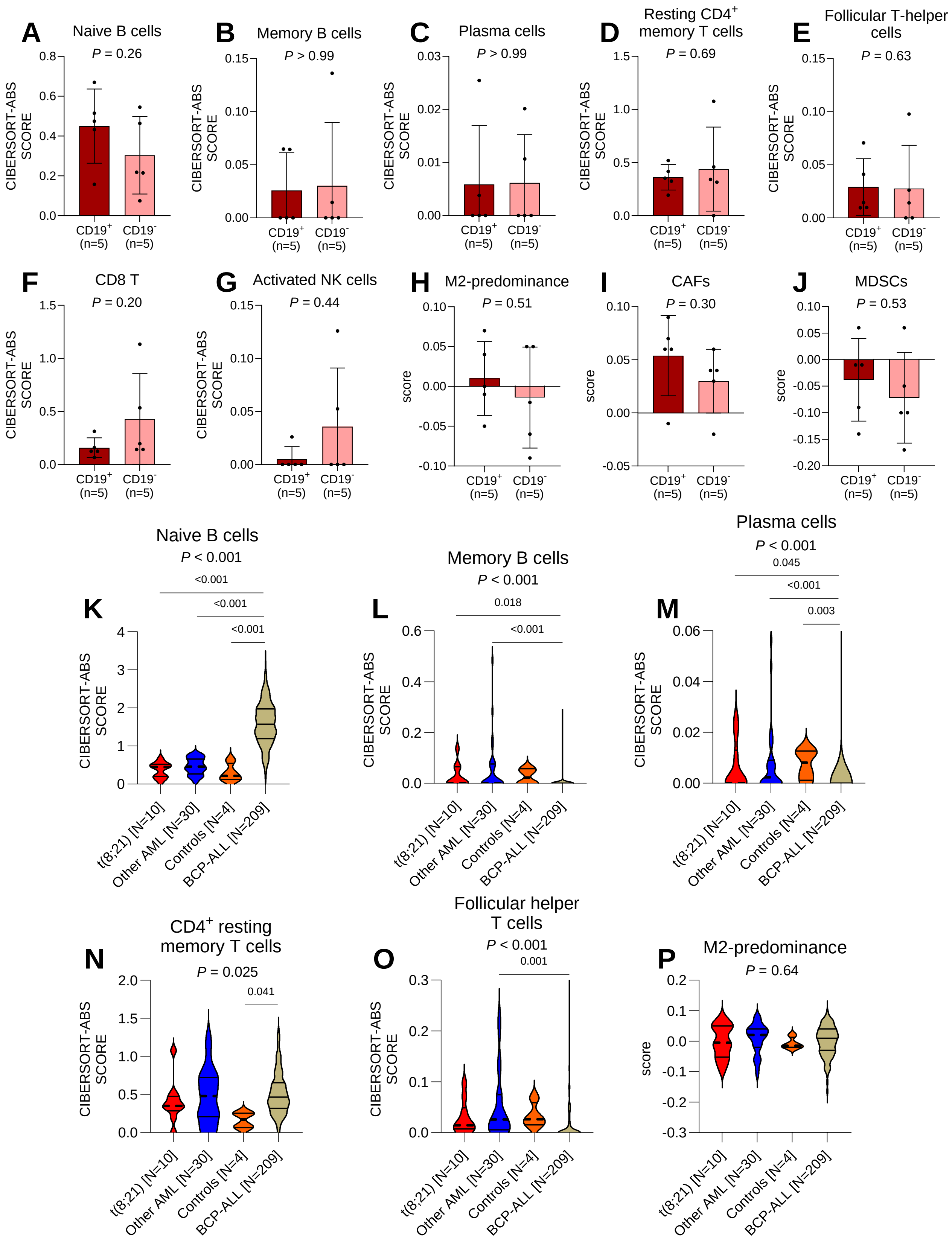
